# Emergence and Enhancement of Ultrasensitivity through Posttranslational Modulation of Protein Stability

**DOI:** 10.1101/2021.08.22.457287

**Authors:** Carla M. Kumbale, Eberhard O. Voit, Qiang Zhang

## Abstract

Signal amplification converts a linear input to a steeply sigmoid output and is central to cellular functions. One canonical signal amplifying motif is zero-order ultrasensitivity through the posttranslational modification (PTM) cycle signaling proteins. The functionality of this signaling motif has been examined conventionally by supposing that the total amount of the protein substrates remains constant. However, covalent modification of signaling proteins often results in changes in their stability, which affects the abundance of the protein substrates. Here we use a mathematical model to explore the signal amplification properties in such scenarios. Our simulations indicate that PTM-induced protein stabilization brings the enzymes closer to saturation, and as a result, ultrasensitivity may emerge or is greatly enhanced, with a steeper sigmoidal response of higher magnitude and generally longer response time. In cases where PTM destabilizes the protein, ultrasensitivity can be regained through changes in the activities of the involved enzymes or from increased protein synthesis. Interestingly, ultrasensitivity is not limited to modified or unmodified protein substrates; the total protein substrate can also exhibit ultrasensitivity. It is conceivable that cells use inducible protein stabilization as a way to boost signal amplification while saving energy by keeping the protein substrate at low basal conditions.

## Introduction

### Regulation of protein stability through posttranslational modifications

It has been known for some while that posttranslational modifications (PTMs) are important mechanisms for regulating not only the activity of a protein, but also the abundance of a protein by means of changing its stability. A well-studied example is the DNA damage response. Once the tumor suppressor p53 is phosphorylated by upstream kinases, such as ATM (ataxia telangiectasia mutated), in response to DNA double-strand breaks, its half-life increases dramatically from less than 30 minutes to over 3 hours (Fig. 1A), which causes the accumulation of p53 that can induce target gene expression [1, 2]. A second example, in some sense of the opposite nature, occurs in the germinal center response of B lymphocytes. B cell receptor-activated MAPK phosphorylates BCL6 (B-cell lymphoma 6), resulting in accelerated degradation of BCL6 by the ubiquitin/proteasome pathway (Fig. 1B), which helps the B cells exit the germinal center response [3]. Many similar examples have been reported where protein stabilization or destabilization drives signaling, including IKK-mediated phosphorylation and degradation of IκB in the inflammatory response, Chk1-mediated phosphorylation and proteasomal degradation of Cdc25A during cell cycle arrest, and stabilization of ΔFosB by casein kinase 2-mediated phosphorylation, which might be responsible for long-term adaptation in the brain [4-6]. It is thus conceivable—and even likely—that altering protein stability and/or activity through the same PTM event may be an important, controllable mode of dual regulation of cellular signaling in general. Expressed differently, if the abundance of a protein substrate can be fine-tuned through changes in protein stability, then these changes can in turn be used by the cell as modulators of both the dynamic and steady-state input-output (I/O) behaviors of covalent modification cycles (CMCs), which may or may not alter the activity of the protein.

**Figure 1.**
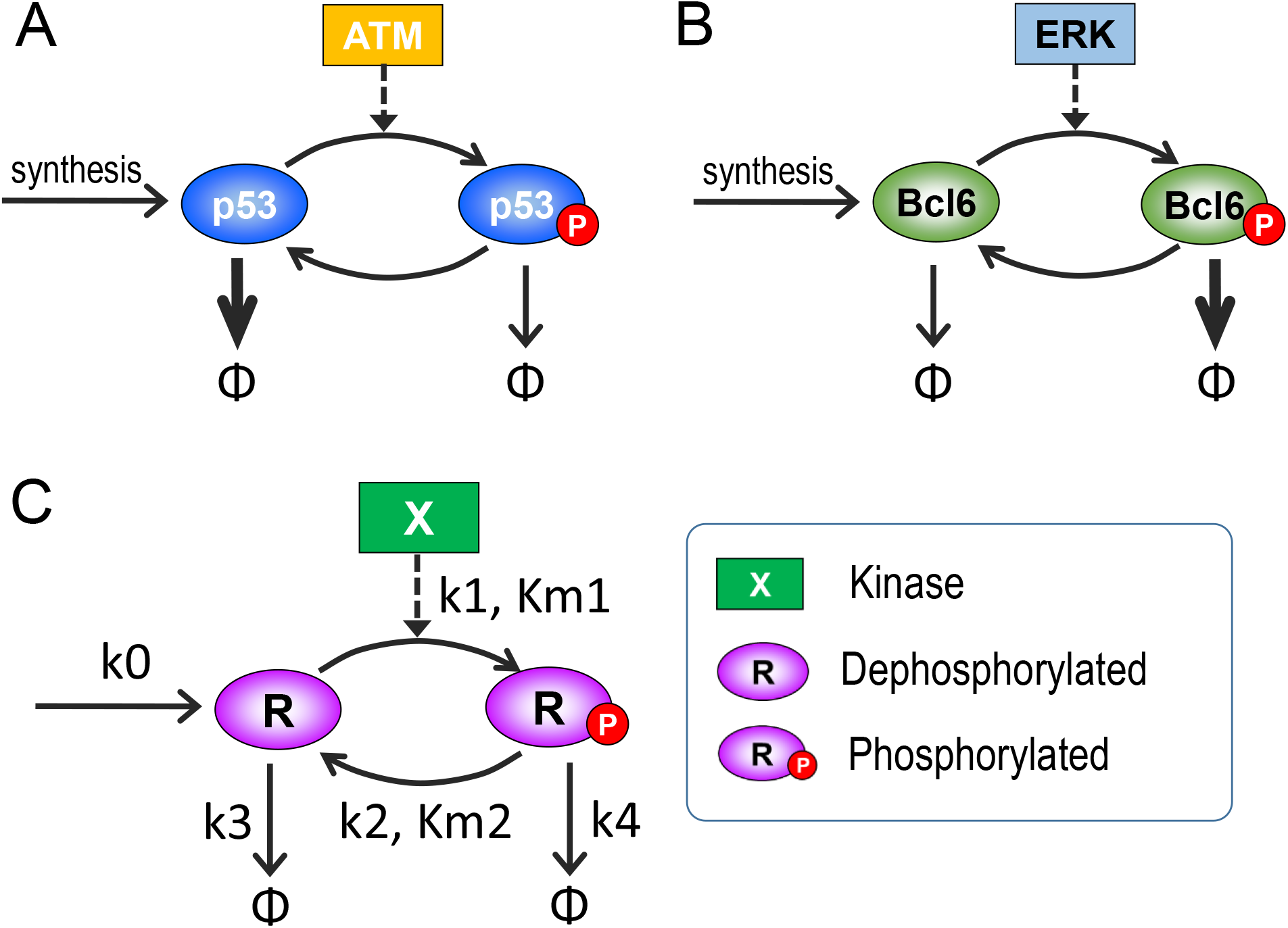
Schematic illustration of covalent protein modification cycles (CMCs) that respond to altered protein stability. (A) p53 stabilization by ATM-catalyzed phosphorylation. (B) BCL6 destabilization by ERK-catalyzed phosphorylation. (C) Generic signaling motif based on phosphorylation-dephosphorylation, used here as the baseline for modeling (*Y*, the phosphatase driving dephosphorylation of *R*_*P*_, is not shown). Open arrow heads: mass flux; thick arrows: fluxes with high degradation rates, dashed arrows: enzymatic catalysis.

### Ultrasensitivity

Cell signaling networks display “ultrasensitivity” if small changes in input are amplified into much larger percentage changes in output [7, 8]. An ultrasensitive I/O relationship is generally sigmoidal in shape and often approximated by a Hill function; the terminology suggests that an ultrasensitive response is steeper than the well-known hyperbolic trend of a Michaelis-Menten function [9, 10]. Embedded in complex network structures such as feedback and feedforward loops, signal amplification is required for cells and organisms to achieve higher-order functions, including differentiation, proliferation, homeostasis, adaptation, and biological rhythms [11, 12]. At least six major ultrasensitive response motifs (URM) have been identified in intracellular molecular networks, namely: (i) positive cooperative binding, (ii) homo-multimerization, (iii) multistep signaling, (iv) molecular titration, (v) zero-order CMCs, and (vi) positive feedback [12-14]. Each of these URMs has its own unique mechanism achieving signal amplification.

### Ultrasensitivity through zero-order covalent modification cycle

The ubiquitous zero-order CMC is particularly interesting, as it can generate nearly switch-like responses. A typical implementation is a modifying / demodifying cycle that is driven by PTMs involving phosphorylation, acetylation, oxidation, methylation, or sumoylation [15]. Specifically, protein activities can be regulated through covalent bonding of moieties to certain amino acid residues, such as phosphate to serine, threonine, and tyrosine in the case of phosphorylation, and an acetyl group to lysine in the case of acetylation. The local electrical charge, possibly accompanied by steric changes introduced by these moieties, can greatly affect the protein molecule’s interaction with other large or small molecules, thereby turning on or off the activity of the protein as an enzyme, transcription factor, or signaling molecule. Covalent modifications of proteins often require specific enzymes, such as kinases, acetyltransferases, methyltransferases, and oxidases, as well as counteracting (demodification) enzymes catalyzing the reverse reactions, such as phosphatases, deacetylases, demethylases, and reductases.

Signal amplification through CMCs was first predicted and analyzed with a mathematical model by Goldbeter and Koshland Jr. in the early 1980s [16, 17]. It occurs when the two opposing enzymes driving the modification cycle of a protein are operating near saturation. In a phosphorylation-dephosphorylation cycle, for example, zero-order ultrasensitivity arises when the amount of protein substrate is at a concentration high enough to saturate the available kinase and phosphatase. Here the terminology “protein substrate” is used to distinguish this protein from the involved enzymes. Under these conditions, small changes in the amount or activity of either the kinase or phosphatase can dramatically change the steady-state fraction of the amounts of phosphorylated or dephosphorylated substrates. Since the theoretical predictions by the Goldbeter-Koshland model, zero-order ultrasensitivity via covalent modification has been reported in numerous biological settings, in both prokaryotes and eukaryotes [18-23].

### Caveat of the Goldbeter-Koshland model suggests a mechanism of signaling control

One important conceptual simplification of the original Goldbeter-Koshland model is that the total abundance of the protein substrate in the CMC is regarded as constant, which ignores turnover via *de novo* protein synthesis and degradation. This omission is possibly critical in the context of protein signaling, as proteins are constantly synthesized and degraded. The assumption of constancy may largely be valid when the signaling events driven by PTM occur rapidly in comparison to the protein substrate turnover. However, even if signaling is fast, it is possible—and indeed a frequent observation as mentioned before—that the PTM alters the stability of the protein substrate, which secondarily affects the total amount of the protein substrate. We first reported that, due to the “leakiness” caused by protein turnover, zero-order ultrasensitivity is compromised when turnover is present, and that the steepness of the sigmoidal response deteriorates as the overall protein turnover rate increases [24]. More recently, Mallela *et al*. further elaborated on the importance of protein synthesis and turnover in affecting zero-order ultrasensitivity of CMCs, especially in the context of multiple PTM cascades sharing the same E3 ligase responsible for protein degradation [25]. Thus, the formerly simple results described by Goldbeter and Koshland are in truth more complicated, as they depend on the kinetic features of the involved enzymes, their saturation, and the degree of protein synthesis and turnover.

Here we pursue the question how cells may use alternate PTM-induced changes in protein stability as an additional layer of control to modulate the zero-order ultrasensitive response of a CMC. In particular, we ask whether such modulations are sufficient to render or enhance ultrasensitivity by stabilizing the protein substrate, or diminish or destroy it by destabilizing the protein substrate. To answer these questions, we systematically study the governing kinetic features of the protein cycle one by one, with mathematical modeling, which allows us to modify any aspect or combination of aspects of a protein signaling cycle with full knowledge of the system features and behaviors. We demonstrate that ultrasensitivity can be gained, enhanced or attenuated for the modified, unmodified, and total protein substrates depending on the conditions of stability changes.

## Methods

### Model structure and parameterization

Our goal is to explore how the behavior of a CMC is affected if protein turnover, protein stability, and kinetic features of the governing enzymes are explicitly taken into account. For this exploration, we consider the generic signaling motif of a protein phosphorylation-dephosphorylation cycle (Fig. 1C) as an “order-of-magnitude” model, *i*.*e*., a numerical model without absolutely precise determination of parameter values and with an expectation of qualitative, rather than quantitative results.

The model consists of two ordinary differential equations (ODEs), formulated in the tradition of mass action and Michaelis-Menten (MM) kinetics:

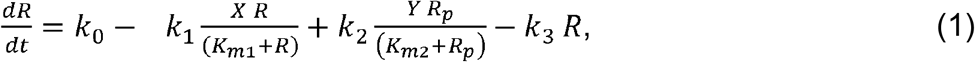

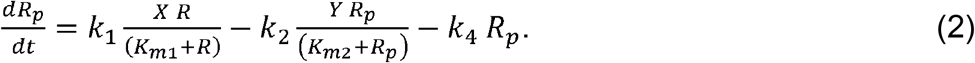

*R* is the protein substrate that is newly synthesized with rate *k*_0_. It can either be phosphorylated into *R*_*p*_ by a kinase *X* which, as a default, follows typical MM kinetics with Michaelis constant *K*_*m*1_ and maximal velocity *V*_*max*1_ = *k*_1_*X*, or it can be degraded with a first-order rate constant *k*_3_. Analogously, *R*_*p*_ can be dephosphorylated by a phosphatase *Y* (not shown in Fig. 1C) that follows MM kinetics with a Michaelis constant *K*_*m*2_ and maximal velocity *V*_*max*2_ = *k*_2_*Y. R*_*p*_ can also be degraded, in this case with a first-order rate constant *k*_4_.

Default parameter values are presented in Table 1. Since covalent protein modifications such as phosphorylation and dephosphorylation occur rapidly, at the order of seconds to minutes, while protein degradation occurs at a much slower rate, often with half-lives at the order of hours, the time scales between these two types of processes are clearly separated by two or more orders of magnitude. Specifically, we set default values for *k*_3_ and *k*_4_ to be 1/100 of *k*_1_/*K*_*m*1_ and *k*_2_/*K*_*m*2_, respectively, because these two ratios approximate the first-order time constants at which phosphorylation and dephosphorylation occur when the kinase and phosphatase are far from saturation. Unless otherwise specified, *Y* is kept as a constant with value 1.

**Table 1.** Default model parameters.

| Parameter | Description | Default Value |
| --- | --- | --- |
| $k_0$ | Rate constant of synthesis of $R$ | 1 (concentration/time) |
| $k_1$ | Catalytic rate constant for phosphorylation | 10 (1/time) |
| $K_{m1}$ | Michaelis constant for phosphorylation | 10 (concentration) |
| $X$ | Kinase | 1 (concentration) |
| $k_2$ | Catalytic rate constant for dephosphorylation | 10 (concentration/time) |
| $K_{m2}$ | Michaelis constant for dephosphorylation | 10 (concentration) |
| $k_3$ | Degradation rate constant of $R$ | 0.01 (1/time) |
| $k_4$ | Degradation rate constant of $R_p$ | 0.01 (1/time) |
| $Y$ | Phosphatase | 1 (concentration) |

### Metrics of ultrasensitivity

In the present study, all dose-response (DR) curves are obtained once the model has achieved steady state. The degree of ultrasensitivity of a steady-state DR curve can be evaluated with two related metrics. First, the Hill coefficient, *n*_*H*_, may be approximated from the equation

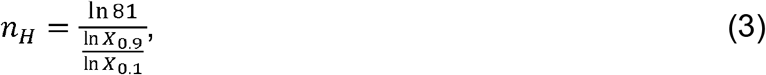

where *X*_0.9_ and *X*_0.1_ are the concentrations of *X* that produce 90% and 10% respectively of the maximal response (after subtracting the background response level when *X*=0) [12]. *n*_*H*_ represents the overall steepness or global degree of ultrasensitivity of the DR curve. Second, we evaluate the local response coefficient (*LRC*) of a DR curve by calculating all slopes of the curve on dual-log scales, which are equivalent to the ratios of the fractional change in response (*R*) to the fractional change in dose (*D*) [7]:

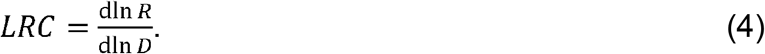

The maximal |*LRC*| of a DR curve (|*LRC*|_*max*_) represents the maximal amplification capacity of the signaling motif. Typical ultrasensitive responses have |*LRC*|_*max*_ values substantially above 1; for values below 1, ultrasensitivity is lost. The comparison between *n*_*H*_ and *LRC* is important as these quantities are not necessarily equivalent and depend on the basal response level and the shape of the DR curve; thus, *n*_*H*_ alone can misrepresent the actual degree of signal amplification [12, 26, 27].

### Simulation tools

The model was coded and simulated in MatLab R2019a (MathWorks, Natick, Massachusetts), which is available as Supplemental Files. All simulations were run using differential equation solver ode23s.

## Results

### 1. Ultrasensitivity in the absence of PTM-induced changes in protein stability

To create a baseline, we start with the default setting *k*_3_=*k*_4_=0.01, which reflects that the phosphorylation status of *R* does not affect its stability. As a consequence, the total steady-state protein substrate concentration *R*_*tot*_ (=*R*+*R*_*p*_) remains constant even if the activity of the kinase *X* varies. Also, the *k*_3_ and *k*_4_ values are very small in comparison to *k*_1_ and *k*_2_. Since *R*_*tot*_ typically exceeds *K*_*m*1_ and *K*_*m*2_ by 10-fold or more, and as expected for the CMC motif, the steady-state DR curves of *R vs. X* and *R*_*p*_ *vs. X* are sigmoidal on the linear scale (Fig. 2A) with *n*_*H*_ at -3.51 and 3.51, respectively (Fig. 2D); the negative sign for *R* indicates a decreasing or inhibitory response. On a log scale, the quasi-exponential rise in *R*_*p*_ and decay of *R* flatten toward straight lines (Fig. 2C).

**Figure 2.**
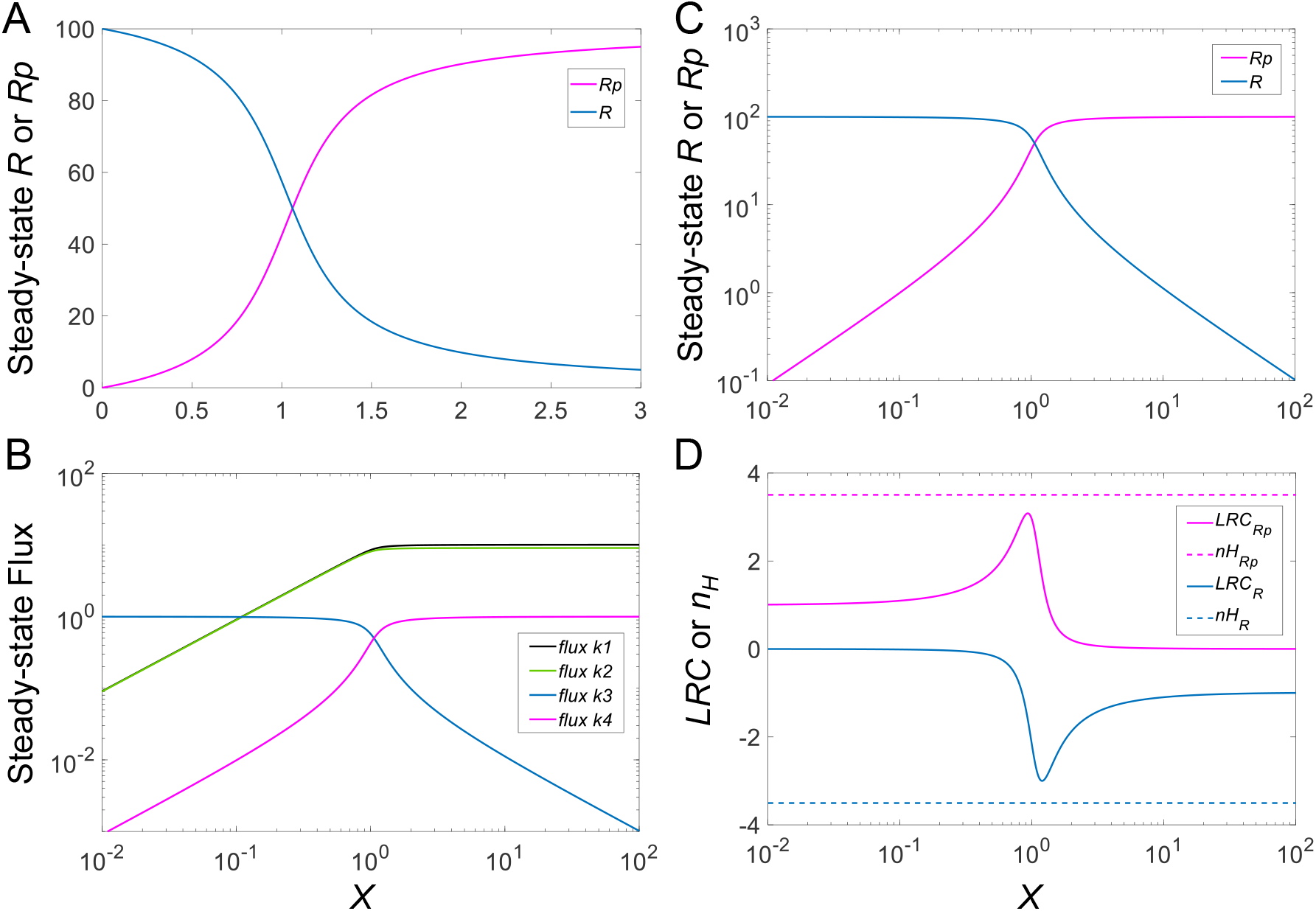
Steady-state DR curves of *R* and *R*_*p*_, associated fluxes, *n*_*H*_ and *LCR*, as functions of kinase activity *X*. **(A)** DR curves of *R vs. X* and *R*_*p*_ *vs. X* on linear scale. **(B)** Fluxes, named by associated rate constant, and plotted against *X*. Specifically, phosphorylation flux: *k*_1_; dephosphorylation flux: *k*_2_; degradation flux of *R*: *k*_3_; and degradation flux of *Rp*: *k*_4_. **(C)** DR curves of *R vs. X* and *R*_*p*_ *vs. X* on double-log scale. **(D)** *n*_*H*_ and *LRC* of DR curves of *R vs. X* and *R*_*p*_ *vs. X*.

The degree of local ultrasensitivity, as measured by *LRC*, varies across the range of *X* and peaks in the center of the DR curves at about -3.0 and 3.1 for *R* and *R*_*p*_, respectively (Fig. 2D). Thus, |*n*_*H*_| in this case is an overestimate of the corresponding |*LRC*|_*max*_. The Phosphorylation and dephosphorylation fluxes (with rates *k*_1_ and *k*_2_, respectively) are dominant over the relatively small protein turnover fluxes with rates *k*_3_ and *k*_4_ at the steady state for large input values of X (Fig. 2B). In a logarithmic representation, these MM fluxes increase essentially linearly as *X* increases before approaching plateaus (Fig. 2B). When protein production and degradation are considered negligible, by setting *k*_0_, *k*_3_ and *k*_4_ to zero, the ultrasensitive responses are slightly enhanced, and the Hill coefficients and (|*LRC*|_*max*_ rise in magnitude to 3.74 and 3.45 for *R*_*p*_ and -3.74 and -3.34 for *R* (simulation results not shown).

### 2. Effects of protein stability on ultrasensitivity

In this section, we suppose that changes in the stability of *R*_*p*_ can be introduced by the PTM, and thus by means of the kinase *X*, as it has been observed numerous times [1, 3-6]. Thus, when the protein substrate *R* is phosphorylated into *R*_*p*_, the stability of *R*_*p*_ is affected, which translates into an increasing or decreasing rate of degradation, *k*_4_. In particular, if the PTM stabilizes *R*_*p*_, *i*.*e*., *k*_4_ decreases, the amount of *R*_*p*_ increases, and *R*_*tot*_ is expected to increase accordingly. This rise in *R*_*tot*_ secondarily alters the degree of saturation of the phosphorylation and dephosphorylation reactions and, consequently, is expected to affect the degree of ultrasensitivity in the DR curves. These overall effects could theoretically also be caused by changes in *k*_3_, but we focus on *k*_4_ because the PTM directly affects the stability of *R*_*p*_, whereas *R* is affected only in a secondary manner.

#### 2.1 Effects on steady-state R

When the PTM increases the stability of *R*_*p*_, *i.e*., *k*_4_ decreases, the steady-state DR curves of *R vs. X* (Fig. 3A) and *R*_*p*_ *vs. X* (Fig. 3B) both become steeper; conversely, when the stability of *R*_*p*_ decreases, *i.e*., *k*_4_ increases, the two curves become shallower. The changes in the steepness of the DR curves can be quantified by *n*_*H*_ and also with the maximal local ultrasensitivity, |*LRC*|_*max*_. Both increase as *k*_4_ decreases (Fig. 3D and 3E). Interestingly, however, for *k*_4_ values comparable to or below the default value, |*LRC*|_*max*_ is generally lower than |*n*_*H*_| for the *R vs. X* response, which is an indication that the Hill coefficient overestimates the maximal degree of signal amplification in these situations (Fig. 3D). For *k*_4_ values rising above the default value, |*LRC*|_*max*_ starts to match up with |n_H_| and eventually even exceeds it. For very large *k*_4_ values, |*n*_*H*_| approaches a constant value of about 1.58 and |*LRC*|_*max*_ approaches a constant value of about 1.72. Thus, there is still ultrasensitivity, but its degree is modest. The value of the kinase activity *X* at which |*LRC*| is maximal shifts to the left as *k*_4_ increases.

**Figure 3.**
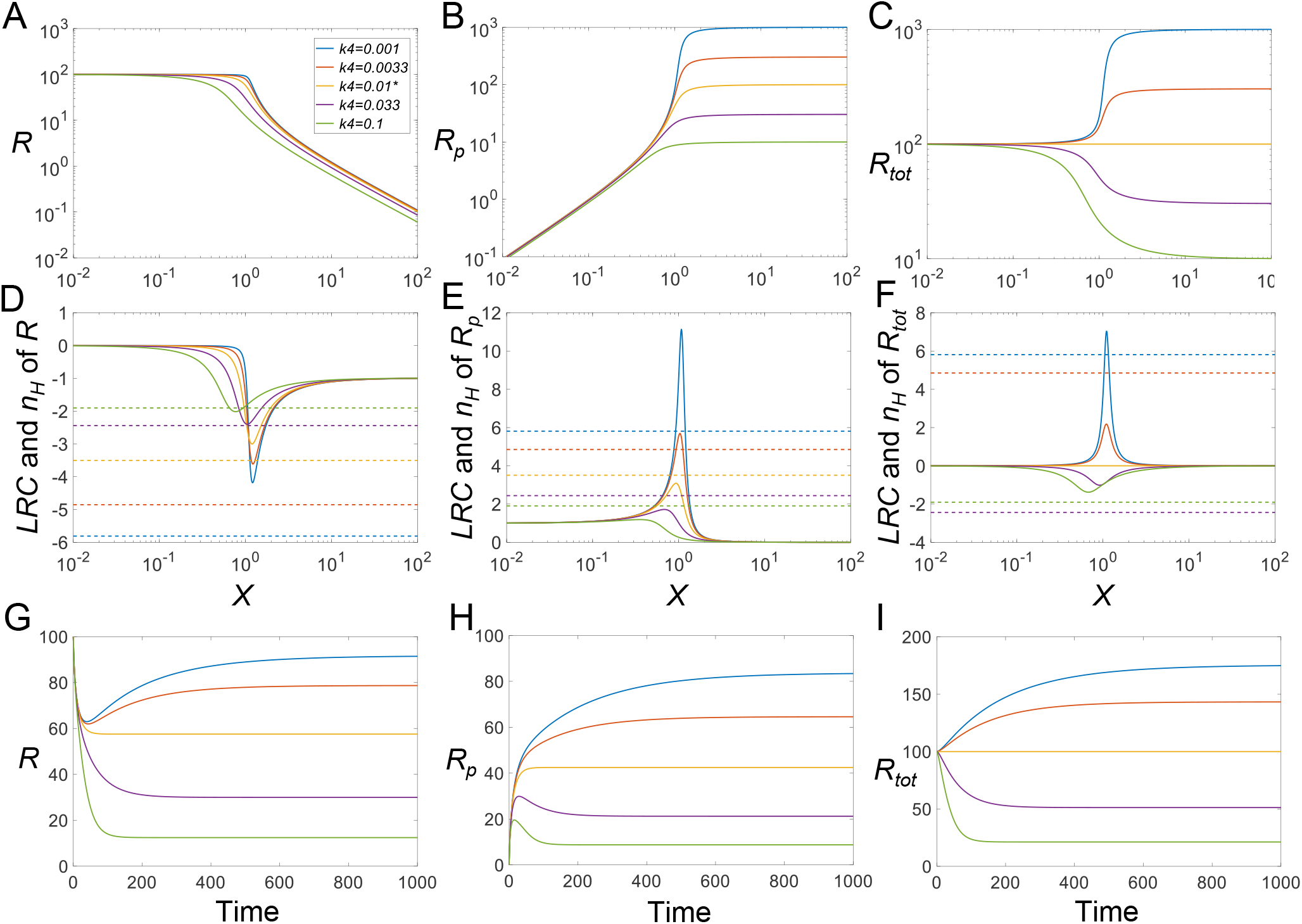
Effects of *k*_4_ on ultrasensitivity and response time. **(A-C)** Steady-state DR curves for *R vs. X, R*_*p*_ *vs. X*, and *R*_*tot*_ *vs. X*, respectively, for different values of *k*_4_, as indicated in panel (A). The color scheme for *k*_4_ in panel (A) is the same for all panels. **(D-F)** *LRC* (solid lines) and *n*_*H*_ (dashed horizontal lines) pertain to *R, R*_*p*_, and *R*_*tot*_, respectively. **(G-I)** Response of *R, R*_*p*_, and *R*_*tot*_ over time, induced by *X*=1, respectively. * *k*_4_=0.01 is the default value.

#### 2.2 Effects on steady-state R_p_

The elevated steepness of the *R*_*p*_ *vs. X* response, with increased stability of *R*_*p*_, is evidently due to the increasing maximal *R*_*p*_ level when *k*_4_ decreases (Fig. 3B). Interestingly, and contrary to the effect on the response of *R vs. X, LRC*_*max*_ is generally higher than *n*_*H*_ for *k*_4_ values below the default value, indicating that the Hill coefficient is underestimating the maximal degree of signal amplification (Fig. 3E). For *k*_4_ values higher than the default value, *LRC*_*max*_ starts to match *n*_*H*_ and eventually drops below its value. For very large *k*_4_ values, *n*_*H*_ approaches 1.58, whereas LRC_max_ settles at about 1. The value of kinase activity *X* for which *LRC* is maximal shifts to the left as *k*_4_ increases.

#### 2.3 Effects on steady-state R_tot_

In the Goldbeter-Koshland model of the CMC, either *R* or *R*_*p*_ is regarded as the output, because the activities of either one may change by the phosphorylation status. However, in some situations, the covalent modification status of an amino acid residue may only affect protein stability without affecting protein activity [28, 29]. In these cases, *R*_*tot*_ should be viewed as the output. Depending on the values of *k*_4_, the response of *R*_*tot*_ vs. *X* can be either stimulatory or inhibitory (Fig. 3C), because either more or less *R*_*p*_ is removed from the system. At the default level of *k*_4_, which is equal to *k*_3_, *R*_*tot*_ does not change with *X*. However, as the PTM stabilizes *R*_*p*_, *i.e*., *k*_4_ decreases from the default value, the steady-state response of *R*_*tot*_ *vs. X* increases monotonically to a higher plateau than before and also becomes increasingly steeper, with *LRC*_*max*_ surpassing *n*_*H*_ for very low *k*_4_ values (Fig. 3F). Conversely, as the PTM destabilizes *R*_*p*_, the steady-state response of *R*_*tot*_ vs. *X* decreases monotonically toward a lower plateau and also becomes increasingly more sigmoidal (despite that the response of *R*_*p*_ itself is no longer ultrasensitive), with |*LRC*|_*max*_ approaching 1.72 for very high *k*_4_ values. Surprisingly, |*n*_*H*_| changes in the opposite direction to |*LRC*|_*max*_ for *k*_4_ values above the default value (Fig. 3F). A small increase in *k*_4_ above the default value first results in a very high |*n*_*H*_|, but as *k*_4_ increases further, |*n*_*H*_| drops back and approaches 1.58. This inverse relationship between |*LRC*|_*max*_ and |*n*_*H*_| demonstrates again that these two metrics do not always conform to each other and that reliance on the Hill coefficient as an estimate of the degree of signal amplification can be misleading. In summary, both stabilization and destabilization of *R*_*p*_ can lead to the enhancement of ultrasensitivity in the steady-state response curve of *R*_*tot*_ *vs. X*.

While *R*_*p*_ and *R* are expected to exhibit ultrasensitivity due to the zero-order covalent modification effect, as revealed by the Goldbeter-Koshland model, it is interesting to note that *R*_*tot*_ also exhibits various degrees of ultrasensitivity depending on the value of *k*_4_, *i.e*., the stability of *R*_*p*_. To dissect this mechanism leading to ultrasensitivity for *R*_*tot*_, we use the following two steady-state flux and mass conservation equations to solve for *R*_*tot*_:

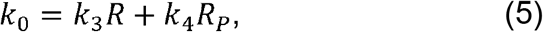

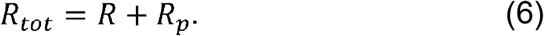

By substituting either *R* or *R*_*p*_ from Eq. (5) in Eq. (6), we obtain two equations that exhibit symmetry with respect to *k*_3_ and *k*_4_, namely

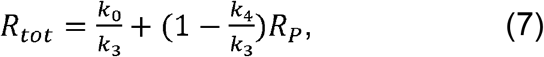

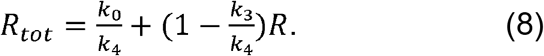

The equations say that except for cases where *k*_3_ and *k*_4_ are equal, the steady-state *R*_*tot*_ scales linearly with both *R*_*p*_ or *R*. When *k*_*3*_>*k*_*4*_, *i.e*., phosphorylation results in *R*_*p*_ stabilization, *R*_*tot*_ has a basal level determined by *k*_*0*_/*k*_*3*_ and increases as *R*_*p*_ increases (Eq 7). For very small *k*_4_, *R*_*tot*_≈*k*_0_/*k*_3_+*R*_*p*_. Since the response curve *R*_*p*_ vs. *X* is always monotonically increasing (Fig. 3B), its ultrasensitivity is passed to *R*_*tot*_ with comparable n_H_ values. By contrast, the LRC of the *R*_*tot*_ response will be lower than that of the *R*_*p*_ response due to the presence of the basal level *k*_0_/*k*_3_ (Fig. 3E *vs*. 3F). Conversely, if phosphorylation results in *R*_*p*_ destabilization, *i.e*., *k*_*3*_<*k*_*4*_, *R*_*tot*_ has a minimal level determined by *k*_0_/*k*_4_ (Eq 8). For very large *k*_*4*_, *R*_*tot*_≈*k*_0_/*k*_4_+*R*. Since the response curve of *R* vs. *X* is always monotonically decreasing (Fig. 3A), its ultrasensitivity is passed to *R*_*tot*_ with comparable *n*_*H*_ values, and again, the |*LRC*| of the *R*_*tot*_ response is lower than that of the *R* response, due to the presence of the minimal level *k*_0_/*k*_4_ (Fig. 3D *vs*. 3F).

#### 2.4 Effects on timing of signaling

PTMs can have an effect on the timing of signaling. When they induce changes in protein stability, the time it takes the signaling motif to reach steady state in response to X is no longer determined only by the covalent modification reactions, but also by the half-lives of the protein substrate. Not surprisingly, for *k*_4_ lower than the default value, it takes much longer time for *R, R*_*p*_ and *R*_*tot*_ to reach their steady state (Fig. 3G-3I). The trajectory of *R* is nonmonotonic – it first decreases quickly as a result of the phosphorylation of pre-existing *R* and then rises slowly (because *R*_*tot*_ increases) to settle at a new steady state (Fig. 3G). In comparison, *R*_*p*_ first shoots up quickly as a result of the phosphorylation of pre-existing *R* into *R*_*p*_, and then rises slowly toward its new steady state (Fig. 3H). *R*_*tot*_ does not exhibit a biphasic trend and instead increases gradually toward its new steady state (Fig. 3I). For *k*_4_ higher than the default value, the time it takes to reach the steady state does not appear to be monotonically correlated with *k*_4_ (Fig. 3G-3I). For *k*_4_ values slightly higher than *k*_3_, the differential stability of *R* and *R*_*p*_ causes the system to approach the steady state slowly because the protein half-life, rather than the fast MM reactions, dominates the long-term kinetics (Fig. 3G and 3H, purple vs. orange lines). But as *k*_4_ increases further, the responses are overall faster since the overall protein half-life becomes shorter (Fig. 3G-3I, green vs purple lines). Generally, *R* first decreases quickly as a result of phosphorylation of pre-existing *R* and then continues to decrease till it settles to a new steady state (Fig. 3G). In comparison, *R*_*p*_ exhibits a nonmonotonic trajectory – it first rises quickly as a result of phosphorylation of pre-existing *R* into *R*_*p*_, and then decreases (because *R*_*tot*_ decreases) slowly to settle at a new steady state (Fig. 3H). *R*_*tot*_ has a similar monotonically decreasing profile as *R* (Fig. 3I).

### 3. Protein stabilization can lead to the emergence of ultrasensitivity

As we demonstrated for a CMC with pre-existing ultrasensitivity, stabilization of *R*_*p*_ can enhance the degree of ultrasensitivity of the responses. In this section, we explore the possibility that stabilization of *R*_*p*_ can render a formerly non-ultrasensitive CMC ultrasensitive. To demonstrate this possibility, we first destroy ultrasensitivity by raising the default values of the Michaelis constants 10-fold, such that *K*_*m*1_=*K*_*m*2_=100. As a result, the cycle no longer exhibits ultrasensitivity for the former default value 0.01 of *k*_4_ (Figs. 4A-4C), as evaluated by |*LRC*|_*max*_ (Figs. 4D-4F). Starting with this new baseline, we now let *k*_4_ decrease below 0.01, which causes *R*_*p*_ to be more stable than *R*. Indeed, the responses, especially the steady-state DR curves for *R*_*p*_ *vs. X* and *R*_*tot*_ *vs. X*, all begin to show a trend toward ultrasensitivity, as the total protein substrate level approaches and eventually surpasses the Michaelis constants *K*_*m*1_ and *K*_*m*2_, thereby pushing the phosphorylation and dephosphorylation cycle toward saturation (Figs. 4A-4C). These results demonstrate that ultrasensitivity can emerge *de novo* with PTM-induced protein stabilization.

**Figure 4.**
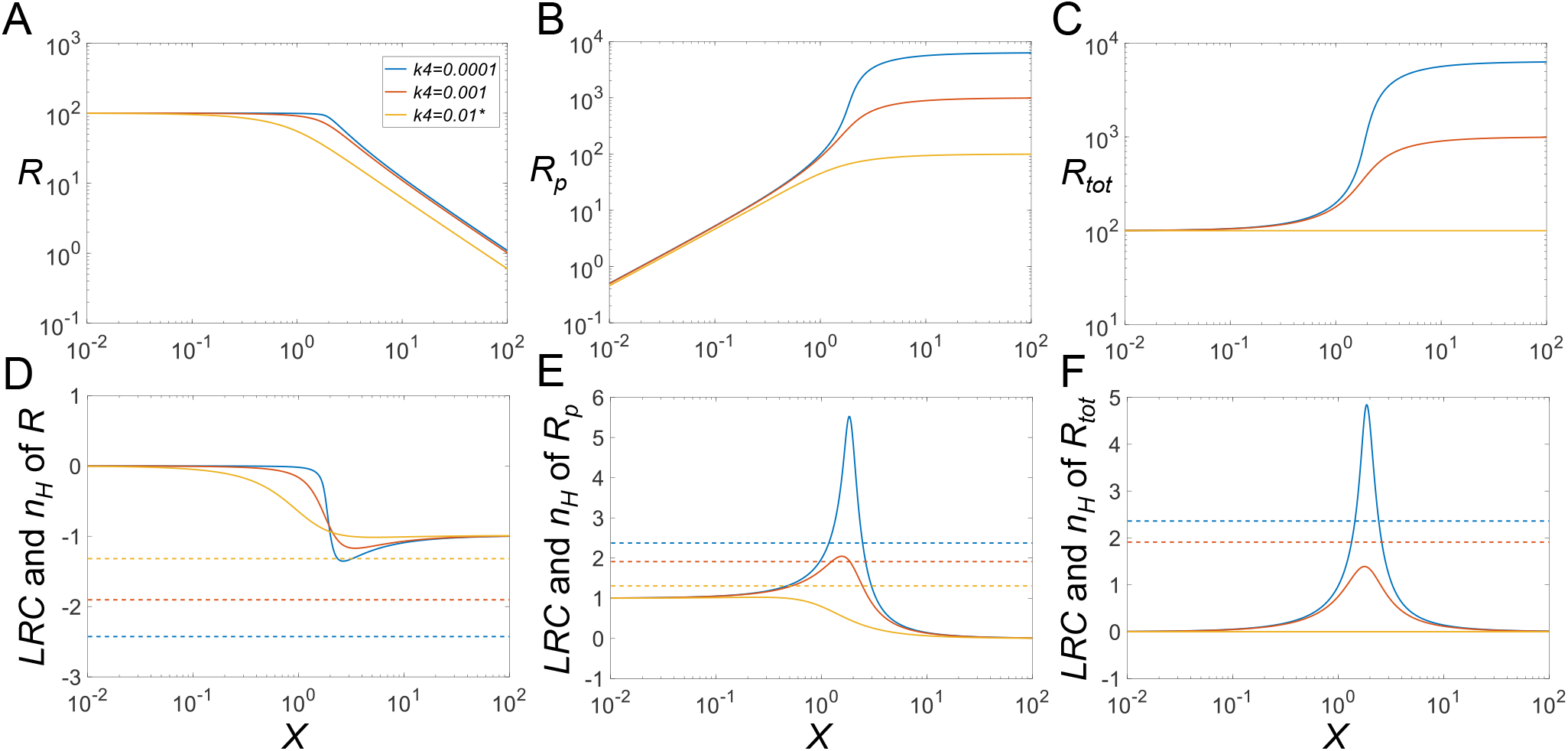
Emergence of ultrasensitivity through phosphorylation-induced protein stabilization. **(A-C)** Steady-state DR curves for *R vs. X, Rp vs. X*, and *R*_*tot*_ *vs. X*, respectively, for different values of *k*_4_, as indicated in panel A. The same color-scheme for *k*_4_ values holds for all panels. As *k*_4_ decreases, ultrasensitivity emerges for *R*_*p*_ and *R*_*tot*_. **(D-F)** *LRC* (solid lines) and *n*_*H*_ (dashed horizontal lines) for *R, Rp*, and *R*_*tot*_, respectively, for different values of *k*_4_. \**k*_4_=0.01 is the default value. For these experiments, the Michaelis constants were set to *K*_*m*1_=*K*_*m*2_=100.

### 4. Regulation of protein modification cycles through alterations in enzyme features

Given the important role of enzyme saturation by the substrate in CMC-mediated ultrasensitivity, we explore in this section whether changes in the kinetic features of the modifying or demodifying enzymes can modulate the DR curves and their ultrasensitivity. Specifically, we investigate how changes in the Michaelis constants *K*_*m*1_ and *K*_*m*2_ modulate the steady-state DR curves and their ultrasensitivity. As a first example, we consider *K*_*m*1_ and examine the case where phosphorylation of *R* into *R*_*p*_ results in destabilization (as the baseline, we set *k*_4_=0.1, which is 10-fold greater than *k*_3_). As *K*_*m*1_ decreases, the DR curves for *R* and *R*_*tot*_ become increasingly more sigmoidal (Figs. 5A and 5C), with limited changes in the *R*_*p*_ responses (Fig. 5B). For low *K*_*m*1_ values, |*LRC*|_*max*_ can be much greater than |*n*_*H*_|, whereas for high *K*_*m*1_ values, |*n*_*H*_| approaches 1.12, and |*LRC*|_*max*_ approaches 1, indicating loss of ultrasensitivity (Figs. 5A and 5D). For the *R*_*p*_ response, increasing *K*_*m*1_ reduces the steepness of the DR curve with |*n*_*H*_| approaching 1.25, and ultrasensitivity is lost for high *K*_*m*1_ values as indicated by |*LRC*| below 1 (Figs. 5B and 5E). Lastly, increasing *K*_*m*1_ reduces the steepness of the DR curve for *R*_*tot*_ with |*n*_*H*_| approaching 1.25, and ultrasensitivity is lost for high *K*_*m*1_ values as is indicated by |*LRC*| below 1 (Figs. 5C and 5F). Varying the Michaelis constant *K*_*m*2_ of the phosphatase has a similar effect on ultrasensitivity (Fig. S1).

**Figure 5.**
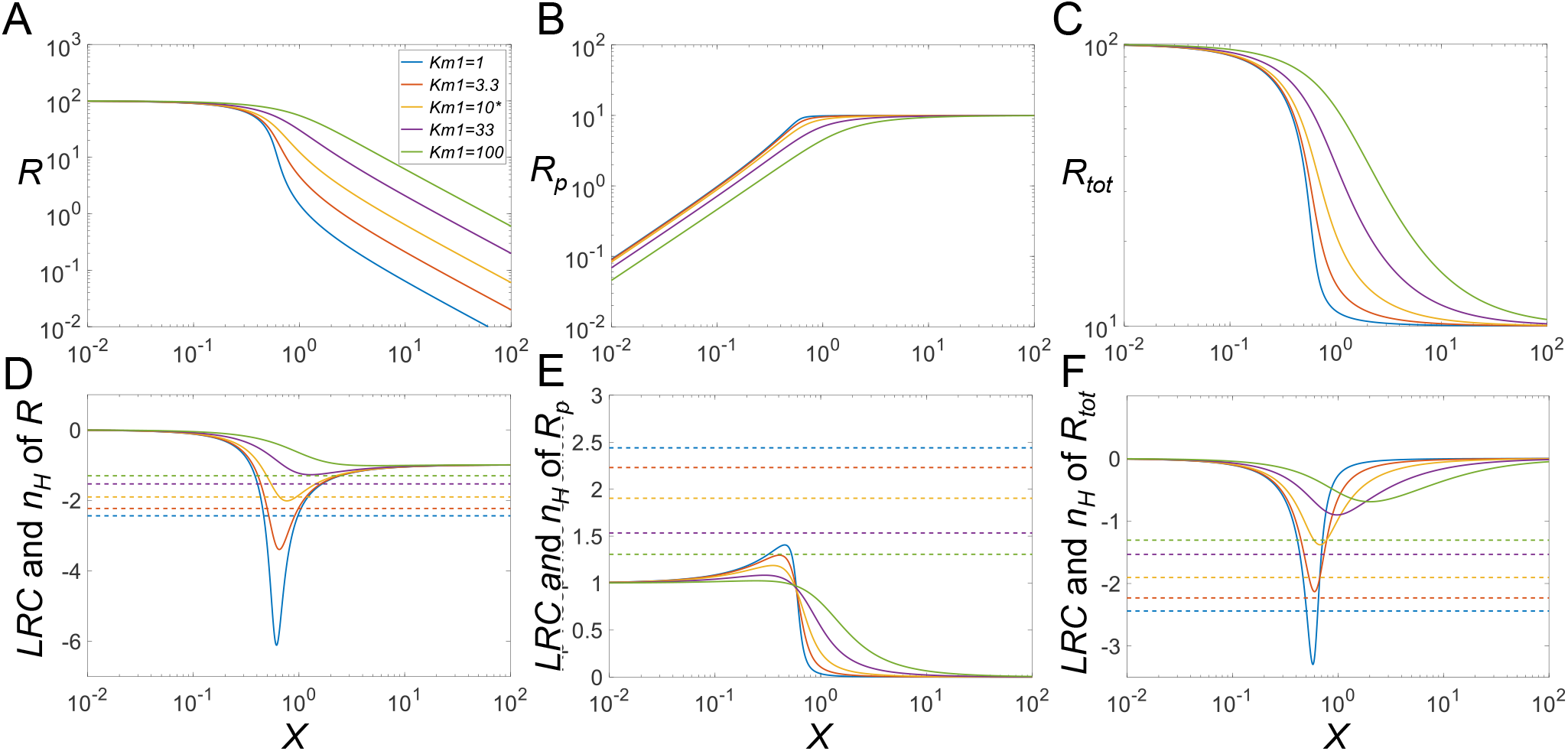
Effects of *K*_*m*1_ on ultrasensitivity under phosphorylation-induced protein destabilization (*k*_4_ = 0.1). **(A-C)** Steady-state DR curves for *R vs. X, R*_*p*_ *vs. X*, and *R*_*tot*_ *vs. X*, respectively, for different values of *K*_*m*1_, as indicated in A. The same color scheme for *K*_*m*1_ values holds for all panels. The degree of ultrasensitivity increases for decreasing values of *K*_*m*1_. **(D-F)** *LRC* (solid lines) and *n*_*H*_ (dashed horizontal lines) for *R, R*_*p*_, and *R*_*tot*_, respectively. * *K*_*m*1_=10 is the default value.

The rationale for a second analysis is the situation where phosphorylation of *R* into *R*_*p*_ results in strong protein stabilization (*k*_4_=0.001, 10-fold lower than *k*_3_). When *K*_*m*1_ decreases below its baseline value of 10 in this situation, the DR curves for *R, R*_*p*_ and *R*_*tot*_ become increasingly sigmoidal. For the response of *R*, |*n*_*H*_| obviously overestimates the degree of ultrasensitivity as evaluated by |*LRC*|_*max*_ (Fig. 6A and 6D). By contrast, for high *K*_*m*1_ values, |*n*_*H*_| approaches 1.93, and |*LRC*|_*max*_ approaches 1, indicating loss of true ultrasensitivity. For the *R*_*p*_ response, increasing *K*_*m*1_ reduces the steepness of the DR curve with |*n*_*H*_| approaching 2.61, and |*LRC*|_*max*_ is reduced to 4.97 with some, but not a complete loss of ultrasensitivity (Fig. 6B and 6E). Except for very high *K*_*m*1_ values, |*LRC*|_*max*_ is generally higher than |*n*_*H*_|. The reason that large *K*_*m*1_ values do not result in complete loss of ultrasensitivity is that *K*_*m*2_ is still kept at default value of 10, thus keeping the dephosphorylation step close to saturable. Lastly, increasing *K*_*m*1_ reduces the steepness of the DR curve for Rp with |*n*_*H*_| approaching 2.61, while |*LRC*|_*max*_ is reduced to 2.34 with some loss of ultrasensitivity (Fig. 6C and 6F). Except for very low *K*_*m*1_ values, |*LRC*|_*max*_ is generally higher than |*n*_*H*_|. Varying *K*_*m*2_ has a similar effect on ultrasensitivity (Fig. S2).

**Figure 6.**
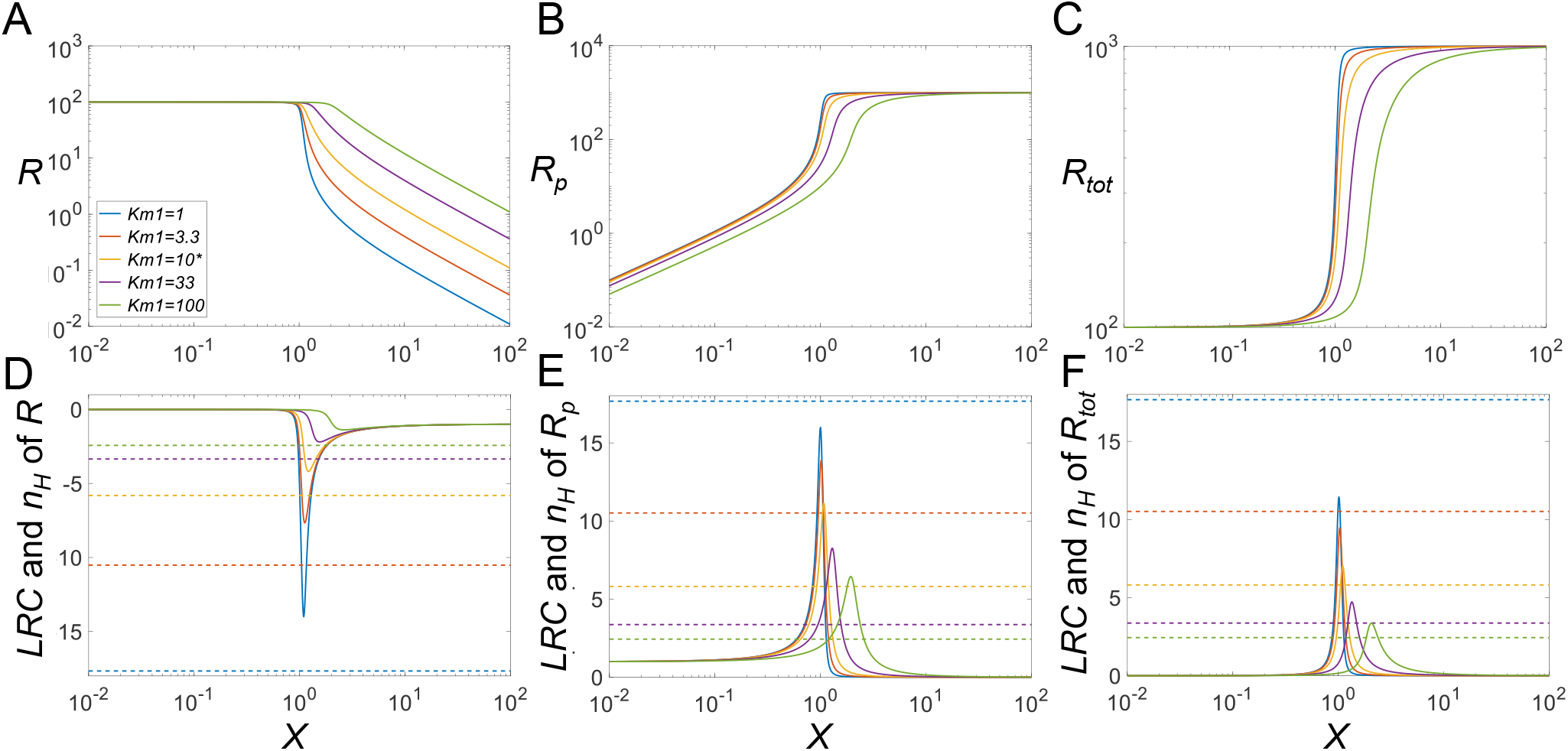
Effects of *K*_*m*1_ on ultrasensitivity under phosphorylation-induced protein stabilization (*k*_4_ = 0.001). In contrast to Figure 5, the results here pertain to a *k*_4_ that is ten-fold lower than the default. **(A-C)** Steady-state DR curves for *R vs. X, R*_*p*_ *vs. X*, and *R*_*tot*_ *vs. X*, respectively, for different values of *K*_*m*1_, as indicated in A. The same color-scheme for *K*_*m*1_ values holds for all panels. **(D-F)** *LRC* (solid lines) and *n*_*H*_ (dashed horizontal lines) for *R, R*_*p*_, and *R*_*tot*_, respectively. * *K*_*m*1_=10 is the default value.

In addition, we studied the effects of changing the catalytic constant *k*_2_ of the phosphatase reaction on ultrasensitivity. In a nutshell, changes in *k*_2_ do affect the degree of ultrasensitivity, but only quantitatively, not qualitatively (Figs. S3 and S4). Varying *k*_1_ merely shifts the DR curves horizontally without changing the degree of ultrasensitivity (simulation results not shown).

### 5. Ultrasensitivity in response to changes in protein synthesis

Lastly, we examine whether changes in the synthesis of *R* can lead to ultrasensitivity if PTM induces changes in protein stability. Suppose the kinase *X* displays an intermediate activity level of 1 and the rate of synthesis of *R, k*_0_, is varied. Interestingly, when *R*_*p*_ is destabilized, *i.e*., *k*_4_> *k*_3_, *R* and *R*_*tot*_ at steady-state exhibit ultrasensitive responses for a certain range of values of *k*_0_ even though their responses never plateau (Fig. 7A and 7C). By contrast, if *k*_0_ is gradually increased, *R*_*p*_ initially increases linearly (in log space), then plateaus, not exhibiting ultrasensitivity for any value of *k*_0_ (Fig. 7B). When *k*_3_=*k*_4_, *R*_*tot*_ is proportional to *k*_0_, and *R* is slightly ultrasensitive. For stabilization of *R*_*p*_, and thus *k*_3_> *k*_4_, the response of *R* vs *k*_0_ is linear, while the response of *R*_*tot*_ vs. *k*_0_ exhibits slight subsensitivity, with *LRC* dipping below 1 for some range of *k*_0_ (Fig. 7F).

**Figure 7.**
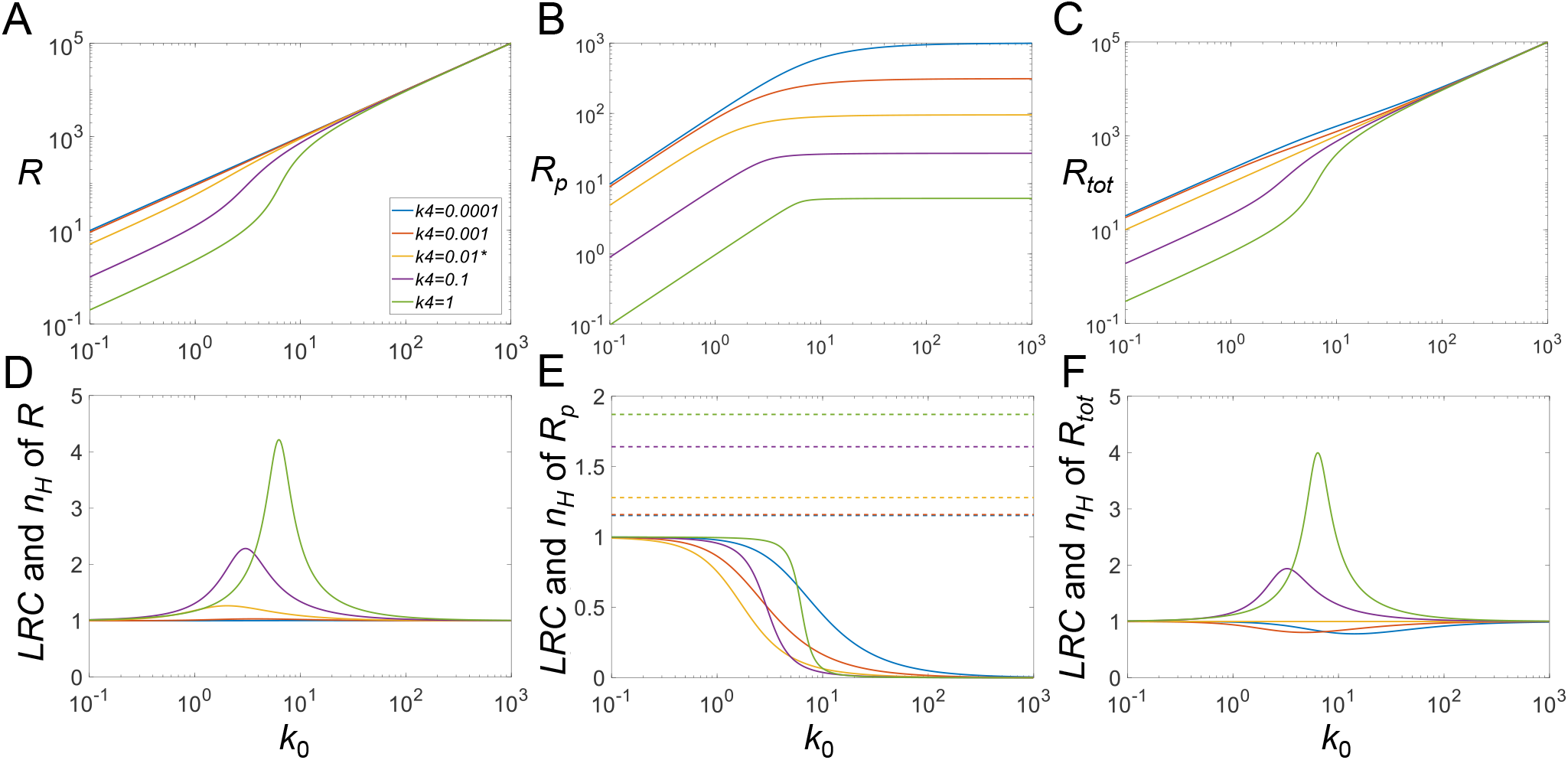
*k*_0_-driven ultrasensitivity with phosphorylation-induced changes in protein stability. **(A-C)** Steady-state DR curves for *R vs. k*_0_, *R*_*p*_ *v.s k*_0_, and *R*_*tot*_ *vs. k*_0_, respectively, for different values of *k*_4_ indicated in (A). The same color-scheme for *k*_4_ values is used for the other panels. **(D-F)** *LRC* (solid lines) and *n*_*H*_ (dashed horizontal lines) for *R, R*_*p*_, and *R*_*tot*_. * *k*_4_=0.01 is the default value. *X*=1 for all conditions. Note that no *n*_*H*_ was evaluated for *R* and *R*_*tot*_ because the responses do not saturate.

The emergence of ultrasensitivity in the responses of *R* and *R*_*tot*_ for high *k*_4_ values may be counterintuitive, since destabilization of *R*_*p*_ is believed to drive the enzymes away from saturation. The reason for ultrasensitivity to occur is the saturation of the flux through the phosphorylation (*k*_1_) step: when *k*_0_ approaches a high value like 10, any further small increase only leads to an increase in *R*, but not *R*_*p*_, and the result is ultrasensitivity. Actually, this mechanism of ultrasensitivity is a variant of zero-order degradation, which no longer requires the dephosphorylation reaction. By setting *k*_2_=0, *i.e*., disabling dephosphorylation, ultrasensitivity in the *R* and *R*_*tot*_ responses remains strong (Fig. S5).

## Discussion

Cellular signal transduction pathways and gene regulatory networks regularly involve PTMs of protein components as a means to regulate their activities and abundance. Nearly all PTM reactions require participation of specific enzymes that add or remove particular functional groups to the appropriate protein substrates. When these enzymes operate near saturation with their substrates, nonlinear signaling may occur, where input signals are amplified to switch output signals on or off [16, 17]. When the protein substrates in a CMC are in excess relative to the modification or demodification enzymes, the degree of saturation of these enzymes depends on the Michaelis constants and the abundance of the contributing substrates.

The covalent modification status of a protein substrate may not only modulate its activity, but also alter its affinity as a substrate for the ubiquitination-proteasomal pathway that mediates the degradation of the majority of intracellular proteins [30]. Depending on whether the covalently modified protein molecule is a better or less suited substrate for ubiquitination, PTM can either stabilize or destabilize the protein and thereby regulate its abundance. For instance, under normoxia, HIF-1α is oxidized by prolyl hydroxylase domain-containing proteins (PHD) in an oxygen-dependent manner and thereby targeted by the pVHL ubiquitination pathway for degradation, thus keeping the hypoxic transcriptional program under control [29, 31]. As a different example, phosphorylation of p53 by ATM during the DNA damage response leads to its stabilization [1]. Therefore, the overall protein half-life and abundance do not remain constant in these situations, rather, they can change dynamically depending on the covalently modified fraction of the protein molecules. The altered protein substrate abundance in turn affects the degree of enzyme saturation, and hence creates an important nonlinearity in signaling.

An obvious scenario of this type is PTM-induced protein stabilization on top of zero-order ultrasensitivity that pre-exists even for basal abundances of the protein substrates. In this scenario, as our simulation demonstrated, the degree of ultrasensitivity for the phosphorylated protein response (*R*_*p*_) with respect to the kinase *X* is considerably elevated, with *LRC* and the Hill coefficient increasing sharply as the half-life of *R*_*p*_ is prolonged (Fig. 3B and 3E). The enhancement of ultrasensitivity is due to the concurrently increased total protein substrate abundance as the input signal *X* increases, which pushes the kinase and phosphatase further into a saturated mode of operation. When the protein substrate is not high enough to enable zero-order ultrasensitivity at the basal condition, the increased protein substrate abundance induced by PTM can move the signaling motif toward saturation, thereby causing the emergence of ultrasensitivity, as demonstrated in Fig. 4B and 4E. During the process of PTM-induced protein stabilization, the unmodified protein response is also enhanced for ultrasensitivity (Fig. 3A, 3D) or rendered ultrasensitive (Fig. 4A and 4D) although the response of *R vs. X* follows an inhibitory profile where *R* decreases as the input signal *X* increases.

An unexpected finding is the total protein response to the input signal (*R*_*tot*_ *vs. X*), which can also exhibit ultrasensitivity, for both cases of PTM-induced protein stabilization and destabilization (Fig. 3C and 4C). The original Goldbeter-Koshland model was intended to examine either the covalently modified or unmodified protein responses under the condition of zero-order ultrasensitivity, while the total protein abundance stayed constant. Here, our simulations show that ultrasensitivity can emerge when there is an imbalance in the stability of the modified and unmodified proteins. When the modified protein is more stable, the total protein response resembles the modified protein response with a non-zero basal level. When modified protein is less stable, the total protein response resembles the unmodified protein response. In both situations, the response of *R*_*tot*_ *vs. X* can be ultrasensitive. As an example, in the drosophila embryo, MAPK can phosphorylate transcriptional repressor Yan in response to morphogen gradients and thereby induce its degradation; this inducible degradation of Yan was proposed as part of a zero-order ultrasensitivity mechanism for the switch-like Yan response responsible for the patterning of the embryonic ventral ectoderm [19]. Therefore, protein activity changes by PTM in a CMC are not mandatory for achieving zero-order ultrasensitivity if protein stability is also regulated by PTM. In the present study, we also demonstrate that if the input-driving signal is supposed to increase the production rate of the protein substrate, a saturable covalent modification reaction, coupled with decreased stability of the modified protein, can also lead to an ultrasensitive increase in either the unmodified or total protein levels (Fig. 7A and 7C).

In the absence of PTM-induced changes in protein stability, the CMC motif can launch a quick response amenable to the time scale associated with covalent modification reactions catalyzed by enzymes. However, when protein stability is altered by PTM with half-lives at the order of hours, it can take much longer for this signaling motif to reach steady state (Fig. 3G-3I). If the protein substrate or its downstream target is a transcription factor, such as p53, HIF-1, BCL-6 or Yan, a relatively slow rise or activation may not matter much as far as the timeliness of a response is concerned, because the ensuing transcriptional induction of downstream genes take much more time to complete anyway. Importantly, we propose here that ultrasensitivity through protein stabilization can be a potential energy-saving strategy employed by cells, where maintaining a high, saturating level of the protein substrate at basal condition may no longer be necessary. In addition, the initial overshoot exhibited by the *R* or *R*_*p*_ response as shown in Fig. 3G and 3I can also be a signaling strategy utilized by cells to accelerate transcriptional induction for gene production with long half-lives [32].

Throughout the result section and the Supplemental Materials, we have compared the degree of steepness of the steady-state DR curve as quantified by *n*_*H*_ with the degree of true ultrasensitivity quantified by *LRC* and confirmed their known differences in describing ultrasensitive DR curves [12, 27]. While the two metrics in most situations move in the same direction in response to changes in a parameter value, the corresponding |*n*_*H*_| for a particular DR curve can be higher or lower than |*LRC*|_*max*_. A higher |*n*_*H*_| value means an overestimate of the degree of amplification of the DR curve, which often occurs when the DR curve has a significant basal level (Fig. 4C and 5C). There are also scenarios where the DR curve exhibits a profile comprising of an almost linear response followed immediately by a plateau (Fig. 7B and S3B). Such a response profile may have an apparent *n*_*H*_=2 despite the fact that its response is at most linear. We have also encountered DR curves having an |*LRC*|_*max*_ value higher than |*n*_*H*_| (Fig. S4B and S4C); in these situations, *n*_*H*_ underestimates the degree of amplification.

Also building upon Goldbeter and Koshland’s concepts, Mallela *et al*. proposed mathematical models for protein modification cycles, focusing, in particular, on protein substrates that are ubiquitinated by the same E3 ligases, which mark both proteins for degradation [25]. Apparently, many E3 ligases are promiscuous, thereby permitting competition between “similar” protein substrates. The authors observed that the sensitivity to incoming signals, as well as the ultrasensitivity of the response, is diminished or even destroyed when the protein substrate saturates the modifying enzyme. This ultrasensitivity-weakening effect is more dramatic if the cycling proteins are degraded at a relatively high rate, consistent with our earlier findings [24]. They also found that signaling cycles, in which the coupling of protein substrates collectively leads to saturation of the enzymes, can lead to a coupled, switch-like response in all protein substrates, likely due to the competition or “crosstalk” of the substrate proteins with respect to the same E3 ligases.

The signaling motif of a CMC can exhibit complex dynamic behaviors and has been extensively studied computationally. Wang *et al*. investigated and decomposed the tunability of the zero-order ultrasensitivity [33]. Xu and Gunawardena examined some more realistic intracellular situations where multiple enzyme intermediates exist due to co-substrate binding for both reversible and irreversible reactions and found that these complications modulate the zero-order switching behavior [34]. The operation of the CMC in the face of protein expression noise has been explored recently [35, 36]. It seems important to have correlated expression of the paired modification and demodification enzymes to prevent switch flipping, and bifunctional enzymes in a CMC may be an ideal solution in this regard [36]. Using linear reactions of the modification and demodification reactions, Soyer demonstrated that the CMC motif, like negative feedback or incoherent feedforward loops, can exhibit transient or persistent dynamic responses depending on the difference in protein stability [37]. As we have demonstrated in the present study (summarized in Table 2), considering PTM-associated changes in protein stability, enzyme features, or protein synthesis can add yet another level of sophistication to the complex response behavior of this long-studied signaling motif.

**Table 2.** Summary of effects of parameters on ultrasensitivity.

| Parameter varied | Parameter condition | Effects on $\vert LRC\vert _{max}$ of DR | | |
| --- | --- | --- | --- | --- |
| | | $R$ vs. $X$ | $R_p$ vs. $X$ | $R_{tot}$ vs. $X$ |
| $\downarrow k_4$ | $k_3 > k_4$ | $\uparrow\uparrow$ | $\uparrow\uparrow$ | $\uparrow\uparrow$ |
| $\uparrow k_4$ | $k_3 < k_4$ | ↓ | ↓ | ↑ |
| $\downarrow K_{m1}$ | $k_3 > k_4$ | $\uparrow\uparrow$ | $\uparrow\uparrow$ | $\uparrow\uparrow$ |
| $\downarrow K_{m1}$ | $k_3 < k_4$ | $\uparrow\uparrow$ | ↑ | $\uparrow\uparrow$ |
| $\downarrow K_{m2}$ | $k_3 > k_4$ | | | |
| $\downarrow K_{m2}$ | $k_3 < k_4$ | $\uparrow\uparrow$ | ↑ | $\uparrow\uparrow$ |
| $\uparrow k_2$ | $k_3 > k_4$ | $\uparrow\uparrow$ | $\uparrow\uparrow$ | $\uparrow\uparrow$ |
| $\uparrow k_2$ | $k_3 < k_4$ | ↑ | ↑ | ↑ |
| | | $R \text{ vs. } k_0$ | $R_p \text{ vs. } k_0$ | $R_{tot} \text{ vs. } k_0$ |
| $\downarrow k_4$ | $k_3 > k_4$ | ↓ | - | ↑ |
| $\uparrow k_4$ | $k_3 < k_4$ | $\uparrow\uparrow$ | - | $\uparrow\uparrow$ |

Note: For the effects on $|LRC|_{max}$ , ↑ or ↓ denotes small effects; $\uparrow\uparrow$ or $\downarrow\downarrow$ denotes large effects; - denotes no effect.

## Acknowledgements

The work was supported by NIEHS Superfund Research grant P42ES04911 and NIEHS HERCULES grant P30ES019776.

