## Supplemental Materials for "Emergence and Enhancement of Ultrasensitivity through Posttranslational Modulation of Protein Stability"

Here we describe the effects of additional means of affecting the regulation of protein modification cycles and ultrasensitivity.

#### *1. Effects of Michaelis constant $K_{m2}$*

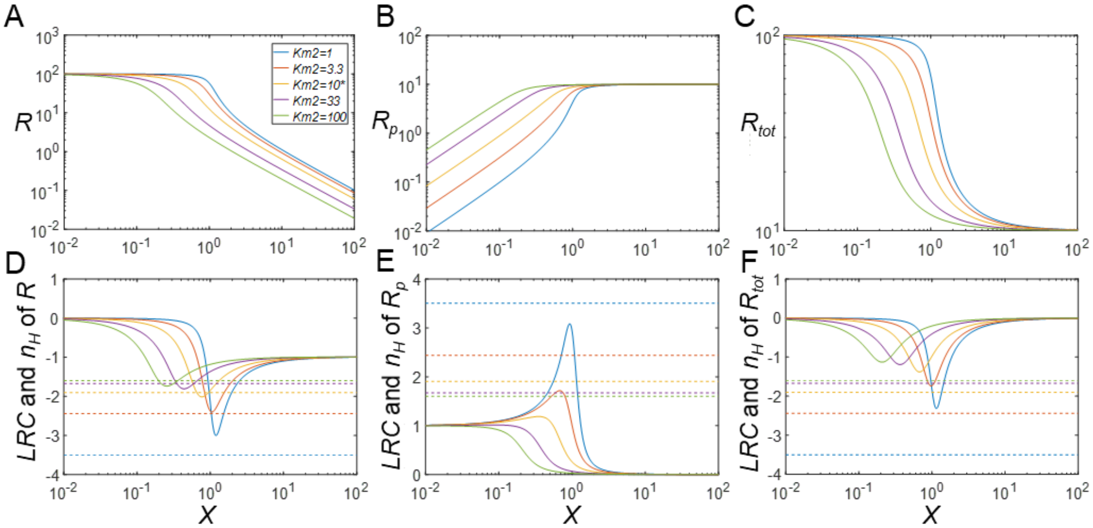


**Figure S1. Effects of $K_{m2}$ on ultrasensitivity with phosphorylation-induced protein destabilization ($k_4$ = 0.1). (A-C)** Steady-state DR curves for $R$ *vs.* $X$, $R_p$ *vs.* $X$, and $R_{tot}$ *vs.* $X$, respectively, for different values of $K_{m2}$, as indicated in panel A. The same color-scheme for $K_{m2}$ is used for all panels. **(D-F)** *LRC* (solid lines) and $n_H$ (dashed horizontal lines) for $R$, $R_p$, and $R_{tot}$, respectively. * $K_{m2}$=10 is the default value.

Here, we explore the role of the Michaelis constant $K_{m2}$ and examine the case where phosphorylation of $R$ into $R_p$ results in destabilization (as the baseline, we set $k_4$=0.1, which is 10-fold greater than $k_3$). As $K_{m2}$ decreases, the DR curves for $R$, $R_p$, and $R_{tot}$ become increasingly steeper (Figs. S1A-S1C). For low $K_{m2}$ values, $|LRC|_{max}$ can be much greater than $|n_H|$, whereas for high $K_{m1}$ values, $|LRC|_{max}$ approaches 1 for $R_p$, indicating loss of ultrasensitivity (Figs. S1B and S1E). For the $R$ response, increasing $K_{m1}$ reduces the steepness of the DR curve without complete loss of ultrasensitivity (Figs. S1A and S1D). Lastly, increasing $K_{m1}$ reduces the steepness of the DR curve for $R_{tot}$ with $|n_H|$ approaching a fixed value slightly below 2, and ultrasensitivity is lost for high $K_{m1}$ values as indicated by $|LRC|$ (Figs. S1C and S1F).

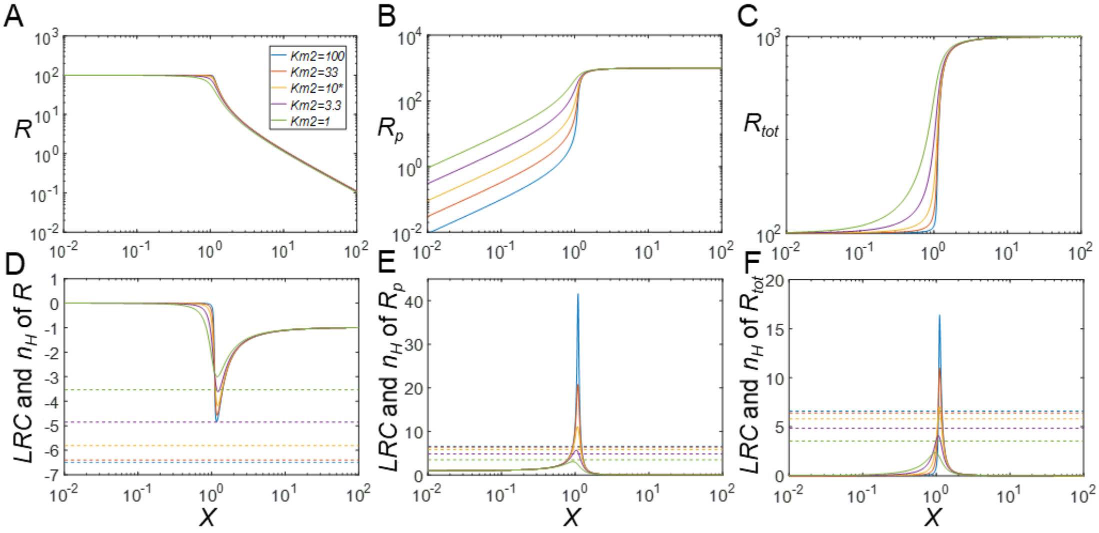


**Figure S2. Effects of $K_{m2}$ on ultrasensitivity with phosphorylation-induced protein stabilization ($k_4$ = 0.001). (A-C)** Steady-state DR curves for $R$ *vs.* $X$, $R_p$ *vs.* $X$, and $R_{tot}$ *vs.* $X$, respectively, for different values of $K_{m2}$, as indicated in panel A. The same color-scheme for $K_{m2}$ is used for all panels. **(D-F)** *LRC* (solid lines) and $n_H$ (dashed horizontal lines) for $R$, $R_p$, and $R_{tot}$, respectively. * $K_{m2}$=10 is the default value.

The rationale for a second analysis is the situation where phosphorylation of $R$ into $R_p$ results in strong protein stabilization ($k_4$=0.001, 10-fold lower than $k_3$). When $K_{m2}$ decreases below its baseline value of 10, the DR curves for $R_p$ and $R_{tot}$ become increasingly steeper with minimal changes in the steepness for $R$ (Fig. S2A-S2C). For the response of $R$, $|n_H|$ obviously overestimates the degree of ultrasensitivity as evaluated by $|LRC|_{max}$ (Fig. S2A and S2D). For the $R_p$ response, increasing $K_{m2}$ reduces the steepness of the DR curve with $|LRC|_{max}$ consistently greater than $|n_H|$ (Fig. S2B and S2E). Lastly, increasing $K_{m1}$ reduces the steepness of the DR curve for $R_p$ with $|n_H|$ overestimating the degree of ultrasensitivity as represented by $|\mathrm{LRC}|_{\mathrm{max}}$ for large $K_{m2}$ values, and underestimating for small $K_{m2}$ values (Fig. S2C and S2F).

### 2. *Effects of catalytic constant $k_2$*

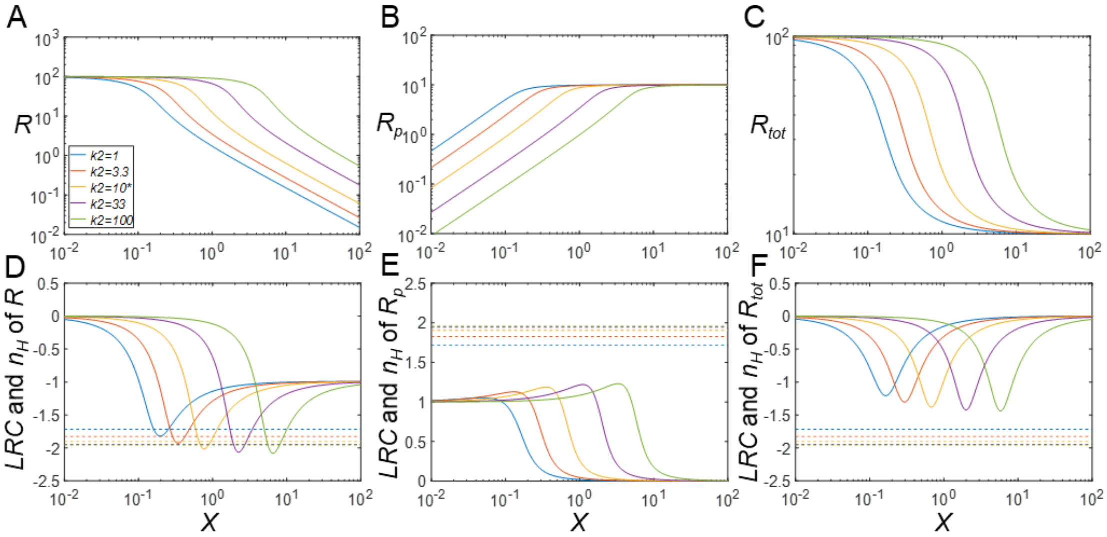


**Figure S3. Effects of $k_2$ on ultrasensitivity with phosphorylation-induced protein destabilization ($k_4$ = 0.1). (A-C)** Steady-state DR curves for $R$ *vs.* $X$, $R_p$ *vs.* $X$, and $R_{tot}$ *vs.* $X$, respectively, for different values of $k_2$, as indicated in panel A. The same color-scheme for $k_2$ is used for all panels. **(D-F)** *LRC* (solid lines) and $n_H$ (dashed horizontal lines) for $R$, $R_p$, and $R_{tot}$, respectively. * $k_2$=10 is the default value.

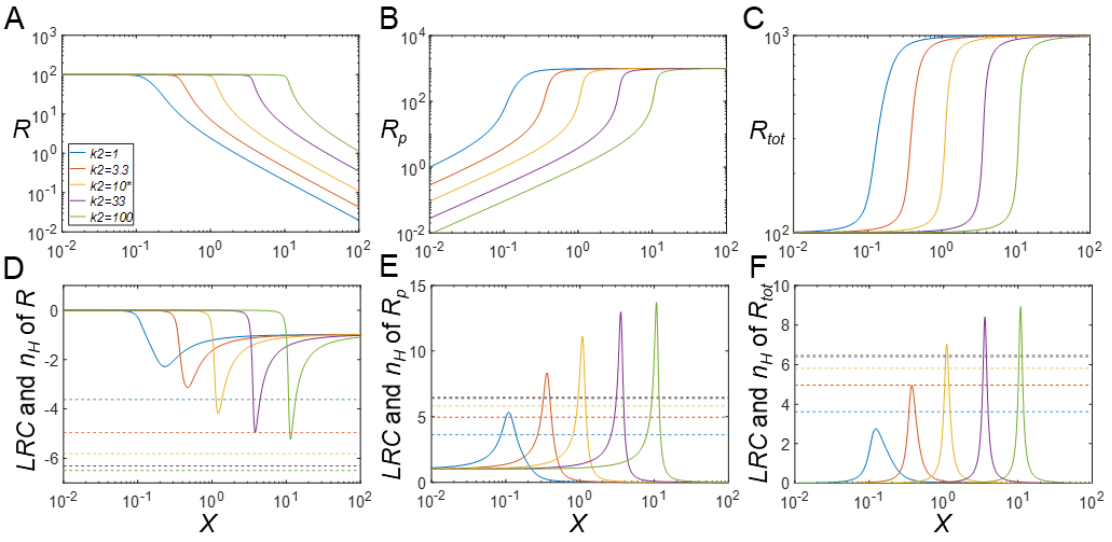


**Figure S4. Effects of $k_2$ on ultrasensitivity with phosphorylation-induced protein stabilization ($k_4$ = 0.001). (A-C)** Steady-state DR curves for $R$ *vs.* $X$, $R_p$ *vs.* $X$, and $R_{tot}$ *vs.* $X$, respectively, for different values of $k_2$, as indicated in panel A. The same color-scheme for $k_2$ values is used for all panels. **(D-F)** *LRC* (solid lines) and $n_H$ (dashed horizontal lines) for $R$, $R_p$, and $R_{tot}$, respectively. * $k_2$=10 is the default value.

Instead of focusing on alterations in $K_m$ values, one may explore the effects of varying $k_2$, the catalytic rate of the dephosphorylation step, on ultrasensitive behavior when changes in protein stability are induced by PTMs. We first examine the case where phosphorylation of $R$ into $R_p$ results in destabilization ($k_4$=0.1, 10-fold greater than $k_3$). As shown in Fig. S3, $k_2$ only has a marginal effect on the degree of ultrasensitivity as assessed by either $n_H$ or *LRC*, while decreasing $k_2$ generally sensitizes all responses, with the DR curves shifting to the left. Increasing $k_2$ appears to increase $|n_H|$ and $|LRC|_{max}$ only slightly, and all DR curves approach a fixed value when $k_2$ becomes very large. For the case where phosphorylation of $R$ into $R_p$ results in protein stabilization ($k_4$=0.001, 10-fold lower than $k_3$), as shown in Fig. S4, increasing $k_2$ desensitizes all responses and shifts the DR curves to the right, thereby increasing the degree of ultrasensitivity. However, for very high $k_2$ values, both $|n_H|$ and $|LRC|_{max}$ seem to reach some maximal values.

### 3. ***Ultrasensitivity in response to changes in protein synthesis***

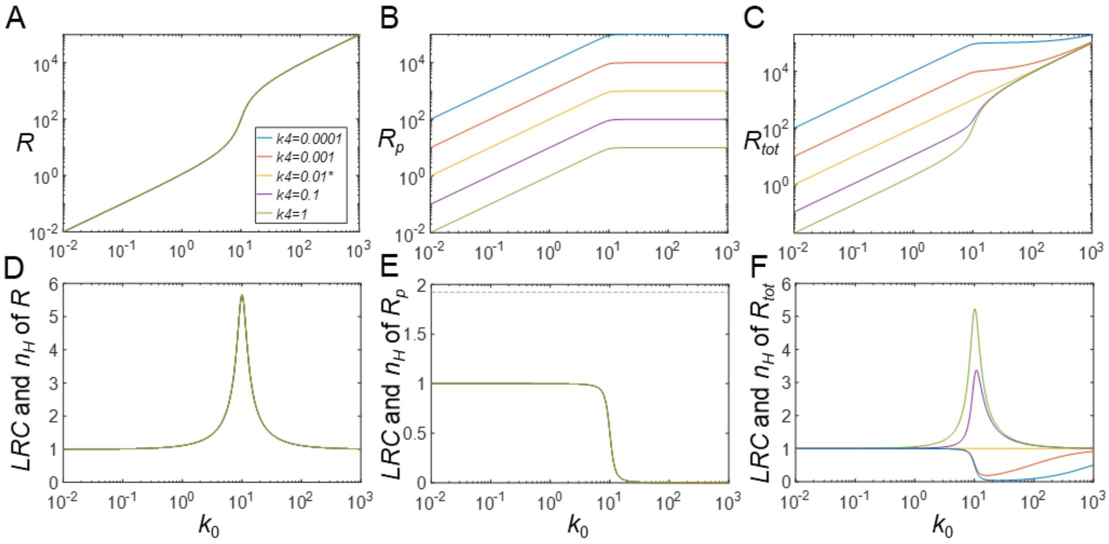


**Figure S5. $k_0$-driven ultrasensitivity with phosphorylation-induced changes in protein stability ($k_2$=0). (A-C)** Steady-state DR curves for $R$ *vs.* $k_0$, $R_p$ *v.s* $k_0$, and $R_{tot}$ *vs.* $k_0$, respectively, for different values of $k_4$ indicated in A. The same color-scheme for $k_4$ values is used for all panels. **(D-F)** *LRC* (solid lines) and $n_H$ (dashed horizontal lines) for $R$, $R_p$, and $R_{tot}$. * $k_4$=0.01 is the default value. $X$=1 for all conditions. Note that no $n_H$ was evaluated for $R$ and $R_{tot}$ because the responses do not saturate. Note: in panels A, D and E, all five curves overlap.

To further explore the mechanism of ultrasensitivity driven by the synthesis rate $k_0$, the dephosphorylation step is disabled by setting $k_2$=0. It is clear for the $R$ vs $k_0$ DR curve, the ultrasensitivity remains regardless the direction in which phosphorylation alters the stability of $R_p$ (Figs. S5A and S5D), suggesting it is the saturation of the phosphorylation that is responsible for the ultrasensitivity. For the $R_{tot}$ vs $k_0$ DR curve, ultrasensitivity is only present when $k_4$>$k_3$, *i.e.*, phosphorylation results in destabilization of $R_p$ (Figs. S5C and S5E), where $R$ dominates the fraction of $R_{tot}$. No ultrasensitivity exists for $R_p$ (Figs. S5C and S5F).
